# Increased biting rate and decreased *Wolbachia* density in irradiated *Aedes* mosquitoes

**DOI:** 10.1101/2021.02.23.432445

**Authors:** Riccardo Moretti, Elena Lampazzi, Claudia Damiani, Giulia Fabbri, Giulia Lombardi, Claudio Pioli, Angiola Desiderio, Aurelio Serrao, Maurizio Calvitti

## Abstract

**Background:** Releasing considerable numbers of radiation-sterilized males is a promising strategy to suppress mosquito vectors. However, releases may also include small percentages of biting females which translate to large numbers when releases are large. Currently, the effects of irradiation on the host-seeking and host-biting behaviors have not been exhaustively investigated. Information is also lacking regarding the effects of sterilizing treatment on the endosymbiotic bacterium *Wolbachia*, which is known to affect the vector competence of infected mosquitos.

**Methods:** To ascertain the effects of irradiation on females, the pupae of two *Aedes albopictus* strains, differing in their natural or artificial *Wolbachia* infection type, and *Ae. aegypti—*which is not infected by *Wolbachia—*were treated with various doses of X-rays and monitored for key fitness parameters and biting behavior over a period of two weeks. The effect of radiation on *Wolbachia* was investigated by qPCR and FISH analysis.

**Results:** Partial *Ae. albopictus* female sterility was achieved at 28 Gy but the number of weekly bites more than doubled compared to that of the controls. Radiation doses of 35 and 45 Gy completely inhibited progeny production but did not significantly affect the survival or flight ability of *Ae. albopictus* females and caused a tripling of the number of bites per female per week (compared to untreated controls). These results were also confirmed in *Ae. aegypti* after treatment at 50 Gy. *Wolbachia* density decreased significantly in 45-Gray-irradiated females, with the greatest decreases in the early irradiation group (26±2-hour-old pupae). *Wolbachia* density also decreased as adults aged. This trend was confirmed in ovaries but not in extra-ovarian tissues. FISH analysis showed a strongly reduced *Wolbachia*-specific fluorescence in the ovaries of 13±1-day-old females.

**Conclusions:** These results suggest that, under SIT programs, the vector capacity of a target population could increase with the frequency of the irradiated females co-released with the sterile males due to an increased biting rate. In the context of a successful suppression, the related safety issues could be generally negligible, but they should be conservatively evaluated when large scale programs relying on imperfect sexing and high overflooding release ratios are run for long time in areas endemic for arboviral diseases. Also, the effects of irradiation on the vector competence deserve further investigation.

## Background

Despite countless attempts, efforts to eliminate vector-borne diseases rarely produce lasting results. In recent years, some vectors have rapidly adapted to the dramatic environmental changes driven by global warming, urbanization, and deforestation, have increased in invasiveness, and have rapidly developed resistance to most pesticides [1–3].

Owing to their reproductive potential and ability to rapidly spread both by migration and passive transportation, mosquitoes are prominent vectors. Substantial investment and research are necessary for the development of prevention and control measures.

The negative effects of widespread insecticide use have caused many researchers to focus on developing innovative control strategies characterized by high specificity and eco-compatibility, and targeting reductions in reproductive potential or vector competence of mosquito populations.

This often involves the release of modified conspecifics who mate with wild types and introduce factors that induce sterility, lethality, or virus resistance in the progeny. To establish such modifications, which may or may not be heritable, advanced biotechnological methods based on genetic modification or the exploitation of beneficial microorganisms are used [4]. Methods based on genetic modification can be targeted to specific genes using various biotechnological tools [5–7] but such approaches can be only employed in countries where the release of GMOs is legal. Sterile Insect Technique (SIT) involves the use of optimized doses of mutagenic radiation to sterilize laboratory-reared males (through non-specific genetic modifications), and releasing large numbers of sterilized males to decrease reproduction in wild populations [8, 9].

A similar result can be obtained via a natural phenomenon of egg inviability induced by the common endosymbiotic bacterium *Wolbachia* [10]. Insect males infected by certain strains of this bacterium are only reproductively compatible with females harboring the same *Wolbachia* strain, whereas the lack of the infection or infection with a non-compatible strain causes the production of inviable offspring. This natural post-mating reproductive barrier is known as Cytoplasmic Incompatibility (CI) and may contribute to the spread of infected individuals in the wild, as infected females can produce viable progeny with both infected and uninfected males (Unidirectional CI, Uni-CI) [11, 12]. The possibility of transferring the bacterium horizontally between species allowed researchers to exploit *Wolbachia* to produce incompatible males that can be released in a target area to reduce the reproductive potential of wild populations (Incompatible Insect Technique, IIT) [13–18].

Large-scale implementation of SIT and IIT must be combined with an efficient method of sexing since the presence of residual females inadvertently released with males can result in the occurrence of temporary spikes in vector density [13]. On the long term, the suppression effect of control programs is expected to counterbalance the contribution of irradiated females to the mean biting rate in the target area. However, even small percentages of co-released females may translate to large numbers of individuals when high overflooding release ratios (10:1 or even 50:1) are applied for long time [19]. Furthermore, especially when *Wolbachia*-induced Uni-CI occurs, released females may be invasive, possibly leading to undesired population replacement [12, 20].

The implementation of SIT using mosquito strains capable of producing incompatible males has been suggested as a method to mitigate the unpredictable effects of female co-release. Mosquito females are more sensitive to radiation than males and this approach could utilize CI to achieve full male sterilization, with the radiation dose lowered to levels that exert no effect on male fitness but which are sufficient to induce complete sterility in females escaping the sexing procedure [21].

However, although recent trials based on this approach have been successful in terms of local *Ae. albopictus* suppression, the strategy has involved the higher radiation doses typical of SIT alone applied on a large scale. This has led to considerably reduced male mating competitiveness without avoiding the release of a few fertile females [19, 22, 23].

It should be noted that all studies aimed at implementing SIT against vector mosquitoes have been mainly focused on radiation doses aimed at the sterilization of the males [9, 24]. However, the radiation doses needed to achieve female sterilization are known to damage the ovarian tissues considerably [21] and previous studies have reported that oogenesis plays a role in the regulation of the tendency to bite and blood-feed [25, 26], which are critical factors for determining the vectorial capacity of mosquitoes [27]. Therefore, an appropriate study on the effects of radiation on these physiological and behavioral traits is warranted.

A thorough investigation on the effects of irradiation on *Wolbachia* would also be necessary in the case of SIT or SIT-IIT combined strategy. In fact, the bacterium *Wolbachia* has been shown to be affected by radiation [28, 29]; thus, this aspect should also be considered before conducting large-scale control programs which involve *Wolbachia* infected mosquitoes as the titer of this endosymbiont may be related to the vector competence of the host [30] and the level of induced CI [31].

Together with certain anopheline species, *Aedes* mosquitoes represent the main concern for human health as serve as vectors for several arboviruses. The impressive spread of diseases associated with these pathogens in recent decades highlights the inefficacy of the current control methods [32–34]. *Ae. albopictus* and *Ae. aegypti* have been the target of several SIT, IIT or SIT/IIT combined experimental trials in recent years [14, 16–19, 35]. The first species is naturally infected with two *Wolbachia* strains that have been demonstrated to interfere with pathogen transmission (compared to the conditions in uninfected individuals) [36] while the latter is not infected by *Wolbachia* in nature.

Herein, we aimed to determine whether irradiation interfered with the host-biting and host-seeking behaviors of *Ae. albopictus* and *Ae. aegypti*. Furthermore, the effects of radiation on the titer of *Wolbachia* in *Ae. albopictus* were evaluated in whole mosquito bodies using qPCR (quantitative polymerase chain reaction), after applying the treatment at different pupal ages and analyzing the effects at two adult ages. Additionally, ovaries and extra-ovarian tissues were similarly studied. Finally, FISH (fluorescence in situ hybridization) analysis was conducted on the irradiated ovaries to visualize the effects of the treatment on the bacterial population and to acquire further information for interpreting the results. Two *Ae. albopictus* lines have been used in the experiments to determine whether different *Wolbachia* strains might affect the results. The resultant data will be useful for enhancing SIT and SIT-IIT strategies in terms of safety and sustainability, and to better evaluate their large-scale applicability.

## Materials and methods

### Ae. albopictus and Ae. aegypti strains and rearing

Two *Ae. albopictus* strains and one *Ae. aegypti* strain were used in the experiments. S_ANG_ *Ae. albopictus* originated from wild-type individuals collected from Anguillara Sabazia (Rome) in 2006 and harbors *w*AlbA and *w*AlbB *Wolbachia*. AR*w*P *Ae. albopictus* was established in 2008 through the transinfection of *Wolbachia*-cured S_ANG_ individuals with *w*Pip *Wolbachia* from *Culex pipiens molestus* [37] and is characterized by a bidirectional incompatibility pattern with wild-type *Ae. albopictus* [38]. Both lines were reared under laboratory conditions at ENEA-Casaccia Research-Center (Rome) and were periodically outcrossed with wild-type individuals from the same area to preserve genetic variability [39]. The *Ae. aegypti* line (New Orleans, LA 2011) was provided by the University of Camerino (Camerino, MC, Italy) where it had been laboratory-reared since 2014 and is not infected with *Wolbachia*.

Colonies were maintained by raising larvae to adulthood inside 1-L larval trays at a density of 1 larva/mL, provided with liquid food according to the methods described in a previous study [39]. Adult mosquitoes were maintained inside 40 *×* 40 *×* 40 cm cages at T = 28 ± 1 C°, RH = 70%±10%, and L:D = 14:10 h, and were supplied with water and 10% sucrose.

Blood meals were provided via anesthetized mice in agreement with the Bioethics Committee for Animal Experimentation in Biomedical Research and in accordance with procedures approved by the ENEA Bioethical Committee according to the EU directive 2010/63/EU. The mice belonged to a colony housed at CR ENEA Casaccia and maintained for experimentation based on the authorization N. 80/2017-PR released (on February 2, 2017) by the Italian Ministry of Health. Feeding female mosquitoes on the blood of human hosts (i.e., the Authors RM, EL, GL, and MC) during the experiments was also approved by the ENEA Bioethical Committee.

### Radiation methods

Cohorts of *Ae. albopictus* and *Ae. aegypti* females belonging to the strains described above were irradiated with X-rays to enable comparison with sham-exposed individuals (control). *Ae. albopictus* pupae were sexed mechanically using a specific sieving procedure described previously [16].

X-ray irradiation was performed using the Gilardoni CHF 320G X-ray generator (Gilardoni S.p.A.; Mandello del Lario, Lecco, Italy) operated at 250 kVp, and 15 mA, with filters of 2.0 mm of Al and 0.5 mm of Cu, furnished by the Physical Technologies for Security and Health Division of ENEA. Depending on the experiment and according to doses already tested for the radiation-based sterilization of the two species [9, 19, 21], *Ae. albopictus* pupae were subjected to 28, 35, and 45 Gy (dose rate: 0.868 ± 0.004 Gy/min, mean ± SD), while a single dose of 50 Gy was used to treat *Ae. aegypti*, as this dosage is known to fully inhibit egg production in the species [9]. Time of sample exposure was determined according to dose rate in order to obtain the pre-established doses. Doses were confirmed by monitoring the exposure with a PTW 7862 large-size plane parallel transmission chamber connected to a PTW IQ4 electrometer. Groups of 100 female pupae were transferred to a Petri dish (d= 4 cm) at 36±4 h of age (unless specified differently) and then transported to the irradiation facility. Immediately before commencement of irradiation, most of the residual water was removed using a glass pipette; irradiated pupae were then transferred to a larger water container to facilitate complete development and allow for adult emergence inside the experimental cages. Sham-exposed pupae, i.e., pupae treated in a manner similar to the exposed pupae except for the X-ray exposure, were used as controls.

### Survival, fecundity, and fertility in irradiated *Ae. albopictus* females

S_ANG_ or AR*w*P *Ae. albopictus* females irradiated at 28, 35, and 45 Gy, and untreated counterparts were allowed to emerge inside 30 × 30 × 30 cm plastic cages, which were checked for the presence of males that escaped the sexing procedure. Thirty virgin females and thirty untreated males— characterized by the same *Wolbachia* infection type—were placed in each cage. These cages were used to monitor survival, fecundity, and fertility in the same treatments during the two weeks of observation, and for the biting rate studies described below. Five repetitions were conducted.

Mortality was recorded daily by removing dead individuals. During the two observation weeks, egg collection was initiated three days after the first blood meal and collection was stopped on the fifth day after the last meal to limit overlap with the second gonotrophic cycle. Paper-lined cups for egg collection were replaced every three days to avoid uncontrolled egg hatching. Eggs were maintained under wet conditions for three days, allowed to dry, and counted to determine the fecundity rate. Egg fertility was assessed by counting the hatched eggs after immersion in a nutrient broth [40].

### Engorgement rate in irradiated *Ae. albopictus* and *Ae. aegypti* females under small enclosures

Starting from the fifth day after emergence and continuing for five days, irradiated S_ANG_ and AR*w*P females were offered a daily blood meal to monitor their feeding behavior. To allow for the study of a second gonotrophic cycle, this same procedure was also conducted from the 12^th^ to the 16^th^ day.

The number of engorged females was tallied within 20 min of placement of the blood meal in the cage. Results obtained with the two populations of females treated at the three radiation doses were compared with those observed using control untreated females. Each of these eight treatments was repeated five times.

Although the present study was focused on *Ae. albopictus*, to generalize the experiment, *Ae. aegypti* females irradiated at 50 Gy were similarly tested (in triplicate) in comparison with untreated controls. The hatch rate of irradiated *Ae. aegypti* was also measured and compared to that of control females to identify a successful radiation treatment.

### Host-seeking ability and biting rate of irradiated *Ae. albopictus* females in large enclosures

The host-seeking behavior of AR*w*P and S_ANG_ females irradiated at 45 Gy was studied in large enclosures and compared with that of untreated individuals based on methods described in a previous study [41]. The trials were conducted outdoor in two large cellular polycarbonate experimental units (LEU, 8.5 × 5 × 5 m L : W : H) with two large lateral openings (L = 8 m; H = 1m) protected by a mosquito net to promote ventilation and ensure natural climatic conditions. The latter were constantly monitored by a CR-10 data logger (Campbell Scientific, Logan, UT, USA) which registered a mean temperature of 34.0 ± 1.0 °C and RH averaging 50.0% ± 5.0%. LEUs contained benches with wet soil and potted plants which provided refugees for the females and higher local levels of humidity (averaging 61.0% ± 5.0%).

For each experiment, an experimenter (the host) as acted as a source of blood and a second experimenter (the collector) released, recovered, and counted the *Ae. albopictus* females after they had fed on the host. The host wore a long-sleeved shirt and short pants, exposing only the lower legs to limit the area for the mosquitoes to land on, whereas the collector wore a white tracksuit and white shoes. Both researchers wore mosquito net hats. Four groups, differing in their infection type (S_ANG_ or AR*w*P) and treatment (irradiated and non-irradiated individuals), were compared. Each group consisted of 30 starved females aged 6±1 days. Females were released at the side of the LEU by removing the cover of the cage. The second experimenter then immediately approached the host on the opposite side of the LEU to collect the females that had started feeding. The proportion of blood-fed females was tallied along with the time taken by each individual to reach the host, at 30-second intervals. Female mosquitoes that landed on the host, but did not bite were excluded from the count. The experiments lasted for a period of 15 min, and for each group, six repetitions were performed, alternating the experimental units and the host. At the end of each experiment, an electric mosquito swatter and a powerful aspirator were used to eliminate any mosquitoes remaining inside the LEU. The experiments were conducted in the late afternoon on sequential days during June 2020. A schematic describing the experiment is provided in the supplemental materials (Additional file 1: Figure S1).

In addition to the methods described above, 6±1-day-old females from each treatment group were engorged in the laboratory and used to study the effect of the radiation treatment on their willingness to seek and bite a host 48 h after the first blood meal. After their release in the LEU, the proportion of feeding females and time to reach the host were again noted.

### Quantitative PCR analysis of *Wolbachia* titer in irradiated *Ae. albopictus* females

Considering the importance of *Wolbachia* in modulating the vector competence of infected mosquitoes [42], a qPCR was performed on the strains of this bacterium present in S_ANG_ and AR*w*P females after irradiation at 45 Gy (a dose known to induce full female sterilization in the species) [9]. Results were compared with those obtained using untreated counterparts. In operational SIT programs, mosquitoes are generally irradiated at the pupal stage (24-48 h); however, it is known that pupal age is one of the critical factors that affects the biological response to the radiation dose [43]. Therefore, the effects of the irradiation were investigated by analyzing DNA extracts of whole bodies of 6±1-day-old females developed from pupae irradiated at three ages (26±2, 36±2 and 46±2 h).

The density of *Wolbachia* is known to vary with the age of *Ae. albopictus* females [44]. Therefore, to investigate the effects of radiation on the bacterium in the germ line and somatic tissues, the titer of the bacterium was measured in ovaries and bodies lacking ovaries of 6±1- and 13±1-day-old females after they were treated with 45 Gy as pupae aged 36±4 h. Untreated females were used as a control.

DNA was extracted from single individuals using the ZR Tissue & Insect DNA Kit MicroPrep (Zymo Research, Irvine, CA, USA), according to manufacturer’s instructions. When necessary, females were chilled on ice and dissected in phosphate-buffered saline (PBS) to isolate the ovaries from the other tissues. Here we excluded individuals with ovaries that were not intact.

Real-time PCR was performed using the Roche LightCycler 96 Instrument (Roche Molecular Systems, Inc., Rotkreuz, Switzerland). Each reaction was performed in triplicate with a μL of the Luna Universal qPCR Master Mix (New England L of 150 nM each primer, and 2 μL of purified DNA) using the following amplification program: initial activation at 95 °C for 120 s, followed by 40 cycles at 95 °C for 15 s and 60 °C for 40 s. The presence of specific amplification products was verified using dissociation curves [44].

Strain-specific primers were used to amplify the *wsp* loci [45], namely, the *w*AlbA-*wsp* and *w*AlbB-*wsp* loci in the case of S_ANG_ females, and the *w*Pip-*wsp* loci in the case of AR*w*P specimens. *Wsp* plasmid standards were used to generate a standard curve [44]. The *Ae. albopictus* actin gene was used as a reference for whole body extracts and extracts obtained after removal of the ovaries and was amplified with the primer pair actAlbqPCR [46]. Owing to the marked decrease in actin gene copies observed preliminarily in irradiated ovaries (Additional file 2: Figure S2), the normalization of qPCR data related to these organs was performed using total DNA (2 μL of purified DNA per reaction, corresponding to 200-300 ng) as reference [47]. For this purpose, NanoDrop 2000 spectrophotometer (Thermo Fisher, Waltham, MA, USA) was used.

### FISH analysis of the ovaries in irradiated *Ae. albopictus* females

Based on the results of qPCR, fluorescent in situ-hybridization (FISH) analysis was conducted on the ovaries of 13±1-day-old irradiated and untreated females according to the protocol described by previous authors [48]. Two hundred nanogram of the *Wolbachia* specific 16S rRNA probe (W2: 5′-CTTCTGTGAGTACCGTCATTATC-3′) was added in the hybridization buffer [43]. Tissues were placed on a slide containing a drop of the VECTASHIELD Antifade Mounting Medium with DAPI (Vector Laboratories, Burlingame, CA, USA) and visualized using the Nikon Eclipse E800 confocal microscope and NIS-Elements 4.0 software (Nikon, Tokyo, Japan). An aposymbiotic population of *Ae. albopictus* [37] was used as the negative control.

## Data analysis

Results were expressed as mean ± SE, and the arcsine square root transformation was applied to analyze proportional data. The Levene’s test and the Shapiro-Wilk test were performed to assess equality of variances and normality, respectively. Statistical analysis was performed using the software PASW statistics (PASW Statistics for Windows, Version 18.0. SPSS Inc., Chicago, IL, USA) with the level of significance set at *P <* 0.05.

The survival curves of the four different treatments for each *Wolbachia* infection type, were compared using the Kaplan-Meier method and the log-rank (Mantel-Cox) test. The Kruskal-Wallis H-test followed by the Conover-Iman test was used to compare fecundity and egg hatch data between treatments within each infection type.

Repeated measures ANOVA (Analysis of Variance) was used to analyze the bite data obtained between treatments over each week. Cages were considered the experimental units and data have been expressed as the mean percentage of engorged females per day, adjusted for cage-specific mortality. If Mauchly’s test indicated a violation of the sphericity assumption, the degrees of freedom were corrected by using Huynh-Feldt estimates. Multiple comparisons between treatments were assessed using Tukey’s HSD post hoc test. Additionally, one-way ANOVA was performed to compare the mean number of bites per female per treatment during each week of study.

Data regarding the host-seeking behavior under large enclosures were analyzed by assigning a value to each mosquito based on the median time of each catching interval to compute the average time to landing. The proportion of engorged mosquitoes was measured based on the ones retrieved within the defined 15-minute interval. Two-way ANOVA was used to analyze the differences between groups in terms of the proportions of females that had bitten the host and to compare average landing times. The Shapiro-Wilk test was conducted to ascertain that the proportions and average landing times were normally distributed.

One-way ANOVA was used to compare within each *Wolbachia* strain, the qPCR data obtained from *Ae. albopictus* females treated at the three tested pupal ages or untreated. The effect of female aging on the overall titer of *Wolbachia* was also analyzed by performing one-way ANOVA within data of each strain of the bacterium. Data regarding ovaries and extra-ovarian tissues of the treated or untreated counterparts were similarly analyzed at the two tested female ages. In the case of rejection of the assumptions of equality of variance and/or normality, the Kruskal-Wallis rank-sum test was performed.

## Results

### Female survival, fecundity, and fertility in irradiated *Ae. albopictus* females

Regardless of the infection type, irradiation slightly reduced the survival of female mosquitoes but this difference was not significant between treatments during the two weeks of observation (Fig 1; Log rank test for S_ANG_: χ^2^ = 2.43; df = 3; P = 0.49; Log rank test for AR*w*P: χ^2^ = 1.847; df = 3; P = 0.61).

**Figure 1.**
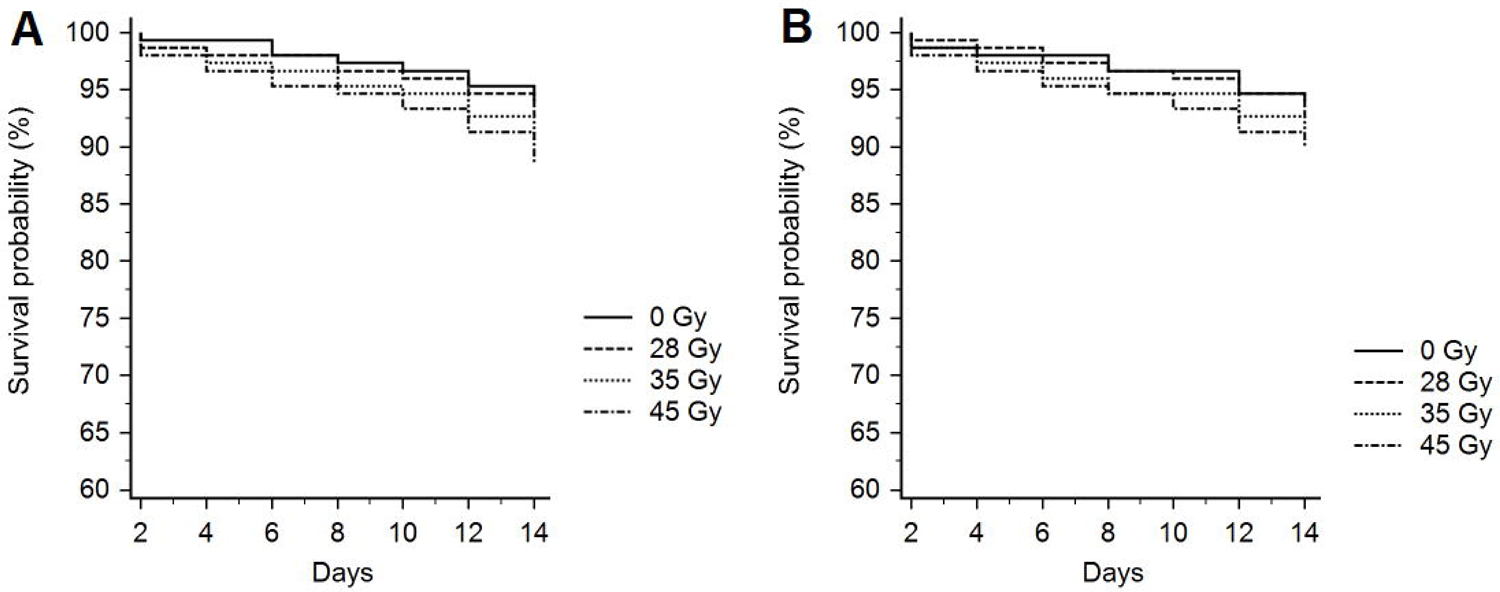
Kaplan-Meier survival curves comparing, respectively, irradiated and untreated S_ANG_ (A) and AR*w*P (B) *Ae. albopictus* over two weeks since the emergence. Differences between treatments were not statistically significant (Log rank test analysis: P < 0.05).

A 45-Gray dose completely inhibited egg laying in *Ae. albopictus*, whereas a few eggs were oviposited by females irradiated at 35 Gy. Among the latter, fertile females were very rare (Table 1). A dose of 28 Gy induced considerable reduction in the number of oviposited eggs compared to that in untreated controls and markedly affected egg fertility, but did not fully inhibit progeny production (Table 1). These results were confirmed in both *Ae. albopictus* strains and both gonotrophic cycles.

**Table 1.**
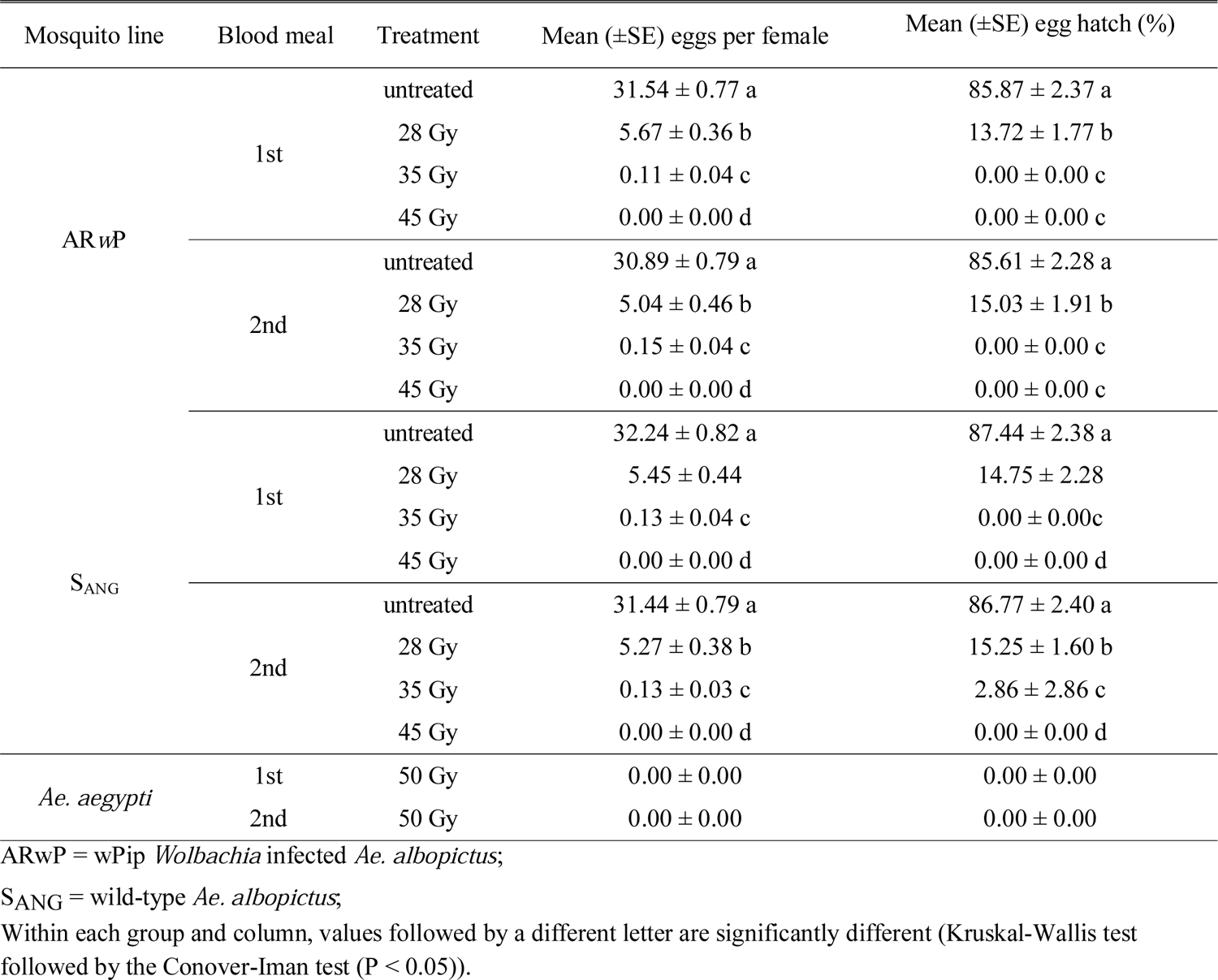
Fecundity and fertility of AR*w*P and S_ANG_ *Ae. albopictus* females irradiated at 28, 35, and 45 Gy compared to those of untreated counterparts over two gonotrophic cycles. Fecundity and egg hatching data related to *Ae. aegypti* irradiated at 50 Gy are also reported.

| Mosquito line | Blood meal | Treatment | Mean (±SE) eggs per female | Mean (±SE) egg hatch (%) |
| --- | --- | --- | --- | --- |
| AR<sub>w</sub>P | 1st | untreated | 31.54 ± 0.77 a | 85.87 ± 2.37 a |
| | | 28 Gy | 5.67 ± 0.36 b | 13.72 ± 1.77 b |
| | | 35 Gy | 0.11 ± 0.04 c | 0.00 ± 0.00 c |
| | | 45 Gy | 0.00 ± 0.00 d | 0.00 ± 0.00 c |
| | 2nd | untreated | 30.89 ± 0.79 a | 85.61 ± 2.28 a |
| | | 28 Gy | 5.04 ± 0.46 b | 15.03 ± 1.91 b |
| | | 35 Gy | 0.15 ± 0.04 c | 0.00 ± 0.00 c |
| | | 45 Gy | 0.00 ± 0.00 d | 0.00 ± 0.00 c |
| S<sub>ANG</sub> | 1st | untreated | 32.24 ± 0.82 a | 87.44 ± 2.38 a |
| | | 28 Gy | 5.45 ± 0.44 | 14.75 ± 2.28 |
| | | 35 Gy | 0.13 ± 0.04 c | 0.00 ± 0.00c |
| | | 45 Gy | 0.00 ± 0.00 d | 0.00 ± 0.00 d |
| | 2nd | untreated | 31.44 ± 0.79 a | 86.77 ± 2.40 a |
| | | 28 Gy | 5.27 ± 0.38 b | 15.25 ± 1.60 b |
| | | 35 Gy | 0.13 ± 0.03 c | 2.86 ± 2.86 c |
| | | 45 Gy | 0.00 ± 0.00 d | 0.00 ± 0.00 d |
| *Ae. aegypti* | 1st | 50 Gy | 0.00 ± 0.00 | 0.00 ± 0.00 |
| | 2nd | 50 Gy | 0.00 ± 0.00 | 0.00 ± 0.00 |

AR<sub>w</sub>P = wPip *Wolbachia* infected *Ae. albopictus*;
S<sub>ANG</sub> = wild-type *Ae. albopictus*;
Within each group and column, values followed by a different letter are significantly different (Kruskal-Wallis test followed by the Conover-Iman test ( $P < 0.05$ )).

### Increased engorgement rate in irradiated *Ae. albopictus* and *Ae*. *aegypti* females

Repeated measures ANOVA revealed significant differences between treatments with respect to the weekly biting activity (Fig. 2A; S_ANG_-1st wk: Huynh-Feldt correction, F_(10.19, 54.35)_ = 2.99, P < 0.05; S_ANG_-2nd wk: Sphericity assumed, F_(12, 64)_ = 4.52, P < 0.05; AR*w*P-1st wk: Sphericity assumed, F_(12,64)_ = 5.90, P < 0.05; AR*w*P-2st wk: Huynh-Feldt correction, F_(9.12, 48.64)_ = 3.36, P < 0.05). Regardless of the week and the infection type, untreated mosquitoes generally showed most biting activity during the first day of the blood meal and then they generally rested while their eggs underwent maturation. In contrast, irradiated females exhibited an anomalous blood feeding behavior and the percentage of the daily engorged females only decreased slightly over each week (Fig. 2A).

**Figure 2.**
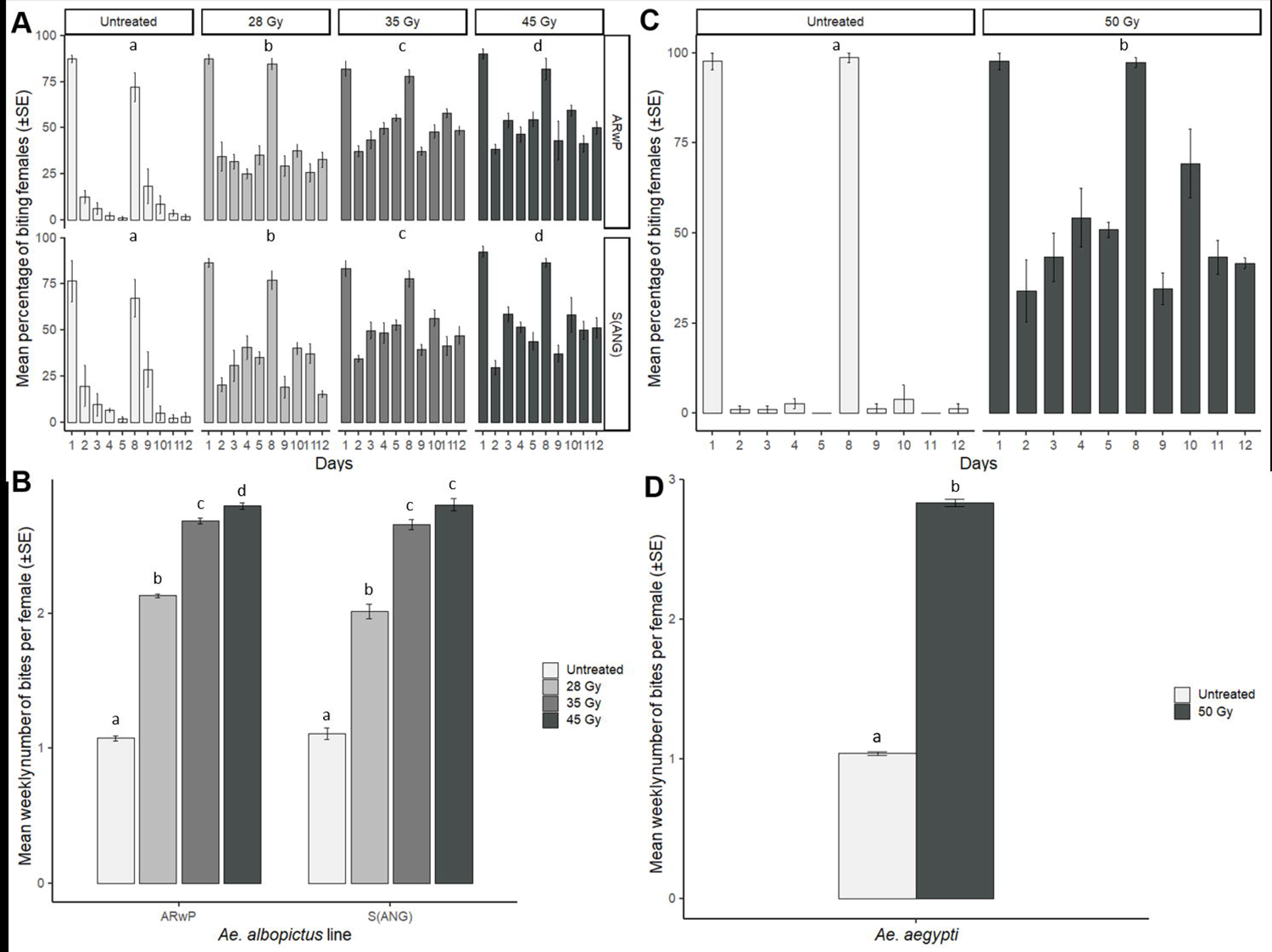
Altered biting behavior in irradiated *Ae. albopictus* and *Ae. aegypti*. The experiment was carried out in laboratory cages by offering daily a blood meal over two sets of 5 subsequent days interrupted by 2 days of rest and starting with 5 days old females. **A:** Daily mean percentage of biting S_ANG_ and AR*w*P *Ae. albopictus*; **B:** Mean weekly number of bites per S_ANG_ and AR*w*P *Ae. albopictus* female over two weeks; **C:** Daily mean percentage of biting *Ae. aegypti*; **D:** Mean weekly number of bites per *Ae. aegypti* female over two weeks; A and C: within each mosquito population, different letters indicate statistically significant differences between treatments (Repeated measures ANOVA followed by Tukey’s HSD test: P < 0.05); B and D: within each mosquito population, different letters indicate statistically significant differences between treatments (One-Way ANOVA followed by Tukey’s test: P < 0.05).

For the 35- and 45-Gray treatments, total bites per female approximately tripled weekly over those recorded for untreated females (Fig. 2B). During the first week, the mean number of bites per S_ANG_ female averaged 1.15 ± 0.07 (untreated), 2.14 ± 0.06 (28 Gy), 2.69 ± 0.07 (35 Gy), and 2.77 ± 0.07 (45 Gy). Tukey’s test showed significant differences among untreated and treated females, and among the females treated with 28 Gy and the other three groups (F_3,16_ = 118.01; *P <* 0.05). During the second week, the above-mentioned values were 1.07 ± 0.05 (untreated), 1.89 ± 0.05 (28 Gy), 2.63 ± 0.03 (35 Gy), and 2.84 ± 0.06 (45 Gy) and all treatments differed significantly from one another (F_3,16_ = 282.23; *P <* 0.05). With respect to the AR*w*P line, the mean number of bites per female was 1.10 ± 0.03 (untreated), 2.15 ± 0.02 (28 Gy), 2.68 ± 0.03 (35 Gy), and 2.84 ± 0.02 (45 Gy) during the first week (F_3,16_ = 371.96; *P <* 0.05), with the Tukey’s test indicating significant differences between all treatments. Values of 1.05 ± 0.03 (untreated), 2.12 ± 0.02 (28 Gy), 2.70 ± 0.03 (35 Gy), and 2.76 ± 0.04 (45 Gy) were obtained during the second week (F_3,16_ = 642.89; *P <* 0.05), with significant differences observed between all treatments except those at 35 and 45 Gy.

Similarly, 50 Gy-irradiated *Ae. aegypti* females showed an enhanced propensity to bite compared to the untreated mosquitoes during both weeks of observation (Fig. 2C; 1st wk: Sphericity assumed, F_(4, 16)_ = 8.38, P < 0.05; 2nd wk: Sphericity assumed, F_(4, 16)_ = 11.51, P < 0.05). Overall, the mean number of bites for the 50-Gray-treated females almost tripled that of untreated mosquitoes during the first (2.81 ± 0.03 and 1.03 ± 0.02; F_1,4_ = 3315.67; P < 0.05) and second weeks (2.86 ± 0.04 and 1.05 ± 0.03; F_1,4_ = 1522.09; P < 0.05) (Fig. 2D). The applied radiation dose was found to induce complete sterility in *Ae. aegypti* females (Table 1).

### Host-seeking behavior of irradiated *Ae. albopictus* females in large enclosures

The host-biting behavior exhibited by the irradiated females in large enclosures reinforced the results obtained in small cages. The applied X-ray dose was not sufficient to reduce the ability of *Ae. albopictus* to reach the host in the LEU (compared to untreated females; Fig. 3A, B). The average times to landing were not significantly affected by the infection type (two-way ANOVA: F_1,20_ = 0.66, P = 0.43), nor were they affected by the treatment (two-way ANOVA: F_1,20_ = 0.93, P = 0.35) in starved females. On average, landing times (±SE) were 346.31 ± 15.98 and 354.30 ± 10.74 in irradiated and untreated S_ANG_ females and 331.11 ± 12.74 and 348.29 ± 12.30 in irradiated and control AR*w*P individuals, respectively.

**Figure 3.**
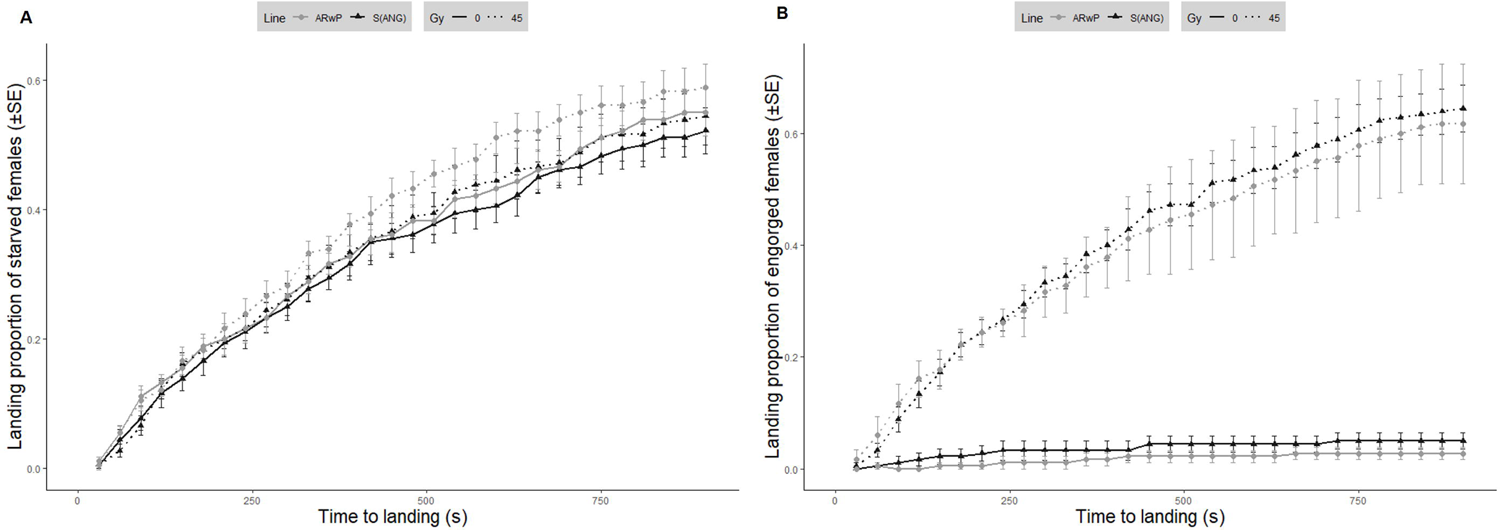
Altered host seeking and biting behavior in irradiated S_ANG_ and AR*w*P *Ae. albopictus* under large enclosures setting. The experiment was carried out in Large Experimental Units (8.5×5×5 m) under open field climatic conditions and involved females irradiated at 45-Gy compared to untreated counterparts. Average landing time and biting proportions were compared between treatments within a 15-minute interval. **A:** Comparison between irradiated and untreated starved females (6±1 days old); **B:** Comparison between irradiated and untreated females 48 h after the engorgement (8±1 days old). Two-way ANOVA demonstrated that the difference between treatments was statistically significant in the case of the engorged females (P < 0.05).

In contrast, the biting behavior was affected by irradiation. Starved females did not reveal a significant difference in the proportion of biting females between untreated and 45-Gray-treated females (two-way ANOVA: F_1,20_ = 0.59, P = 0.45) or between infection types (two-way ANOVA: F_1,20_ = 0.80, P = 0.38) (Fig. 3A). Instead, the proportion of fed females that repeated the blood meal after 48 h was significantly higher in irradiated mosquitoes than that in untreated mosquitoes (two-way ANOVA: F_1,20_ = 276.53, P < 0.05), while the infection type showed no significant effect (two-way ANOVA: F_1,20_ = 1.13 P = 0.30) (Fig. 3B). Consequently, a comparison between average landing times related to the engorged females was not performed due to the reduced number of biting individuals among the untreated controls (0 in certain repetitions). To verify this result in older females that might have experienced the occurrence of age-related damage due to irradiation, the test was also repeated in triplicate with treated and untreated individuals aged 13±1 days and similar results were obtained (Additional file 3: Figure S3).

### Decreased titer of *Wolbachia* in irradiated *Ae. albopictus* females

The age at which *Ae. albopictus* pupae had been irradiated at 45 Gy was found to significantly affect the density of *Wolbachia* (*w*AlbA: F_(3, 36)_=2.96, P < 0.05; *w*AlbB: F_(3, 36)_=4.05, P < 0.05; *w*Pip: F_(3, 36)_=6.76, P < 0.05) regardless of the *Wolbachia* infection type. An early irradiation (26±2-hour-old pupae) resulted in significantly decreased the whole-body *Wolbachia* titer, by a third in *w*AlbA, and by half in *w*AlbB and *w*Pip strains (Fig. 4A) compared to the levels observed in the controls. The *Wolbachia* titer did not significantly differ in pairwise comparisons between treated and untreated individuals when irradiation was performed on older pupae (36 and 46±2-hours old).

**Figure 4.**
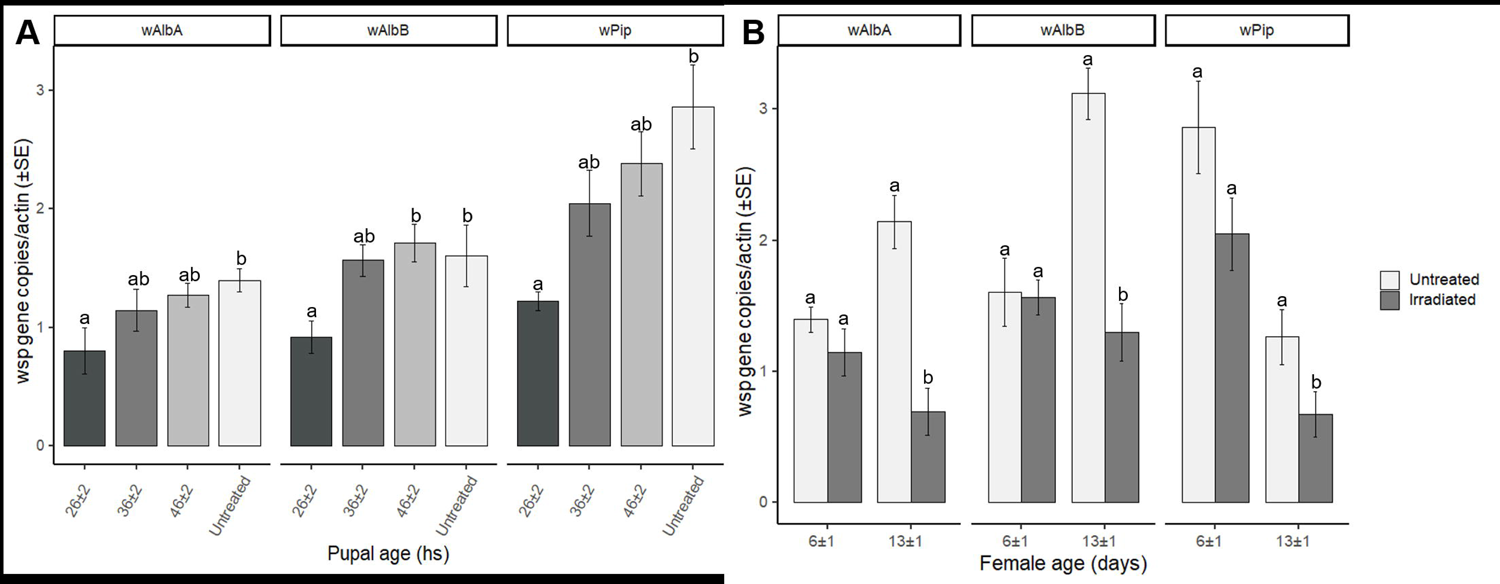
Decreased *Wolbachia* titer in irradiated S_ANG_ and AR*w*P *Ae. albopictus.* **A:** A 45-Gy dose was applied at three different pupal ages (26, 36, and 46±2 hours); quantitative PCR analysis was then performed by using primers targeting the specific *Wolbachia* strains characterizing the two *Ae. albopictus* populations and analyzing 6±1 days old females. **B**: A 45-Gy dose was applied to pupae aged 36±4 hours; qPCR analysis was then carried out studying two female ages (6 and 13±1 days). Data were normalized by using the actin gene as host reference. Within each infection type and female age, different letters indicate statistically significant differences between treatments (One-way ANOVA followed by Tukey’s test: P<0.05).

Aging of adult females was also found to affect the titer of all the tested *Wolbachia* strains when a 45-Gray treatment was performed (Fig. 4B). The difference between treatments was not significant when the analysis had been performed on 6±1-day-old females (one-way ANOVA; *w*AlbA: F_(1, 18)_ = 1.54, P = 0.23; *w*AlbB: F_(1, 18)_ = 0.82, P = 0.89; *w*Pip: F_(1, 18)_ = 3.28, P = 0.09), but the differences increased significantly in females aged 13±1 days (one-way ANOVA; *w*AlbA: F_(1, 18)_ = 62.11, P < 0.05; *w*AlbB: F_(1, 18)_=38.93, P < 0.05; *w*Pip: F_(1, 18)_=4.60, P < 0.05).

In agreement with the data obtained from the analysis of whole bodies, qPCR revealed a correlation between aging and a reduced titer of the bacteria in the ovaries of irradiated *Ae. albopictus* females compared to that in untreated *Ae. albopictus* females (Fig. 5A). Differences were significant in females at 13±1 days after the irradiation (one-way ANOVA; *w*AlbA: F_(1, 18)_ = 62.11, P < 0.05; *w*AlbB: F_(1, 18)_ = 5.91, P < 0.05; *w*Pip: F_(1, 18)_ = 4.60, P < 0.05), whereas the density of *w*AlbA and *w*AlbB *Wolbachia* strains did not differ between treatments in 6±1 day-old females (one-way ANOVA; *w*AlbA: F_(1, 18)_ = 0.39, P = 0.54; *w*AlbB: F_(1, 18)_ = 0.50, P = 0.49). A significant difference was found between the *w*Pip *Wolbachia* titer in treated and untreated 6±1-day-old AR*w*P females (F_(1, 18)_ = 5.25, P < 0.05). The titer of *w*AlbA and *w*AlbB *Wolbachia* also increased with aging in the ovaries of control females and confirmed the results of previous studies (Fig. 5A) [40]. With respect to *w*Pip, the controls exhibited an age-dependent decrease in titer and irradiation further intensified this trend in the density of the bacteria.

**Figure 5.**
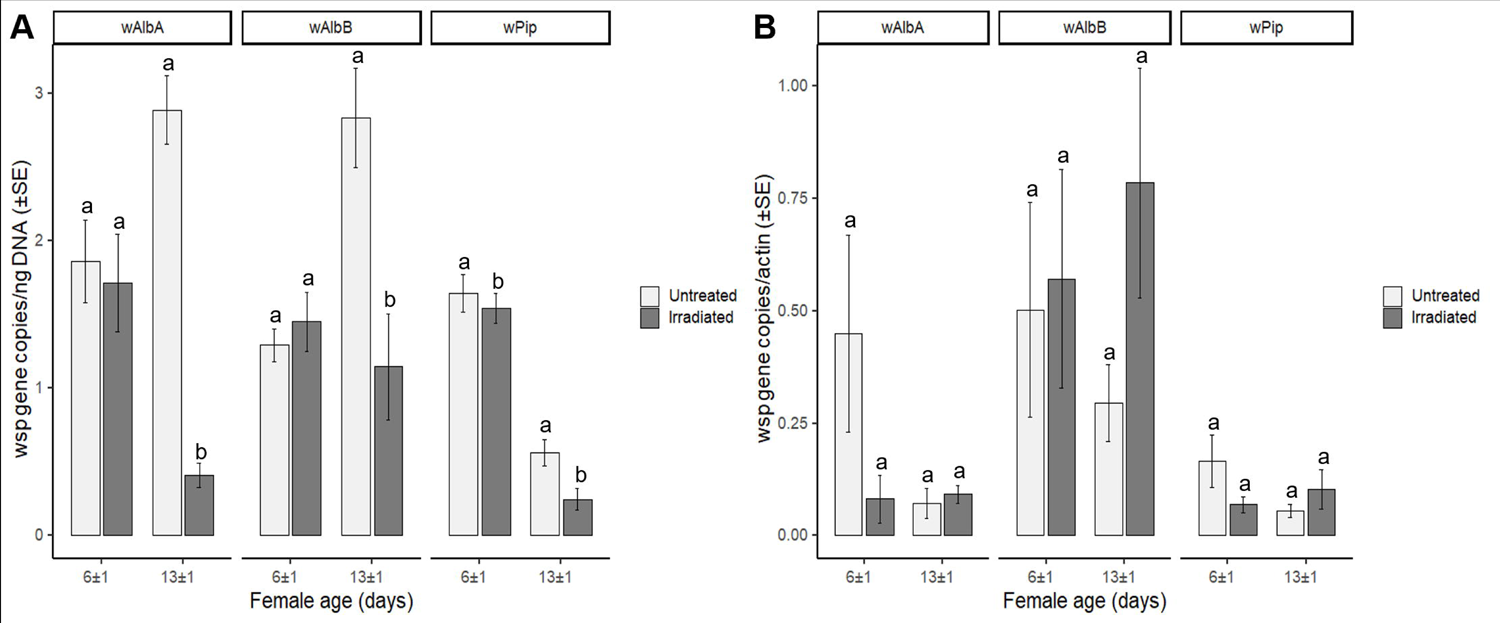
Decreased *Wolbachia* titer in the ovaries of irradiated S_ANG_ and AR*w*P *Ae. albopictus.* A 45-Gy dose was applied to 36±4 hours old pupae. **A:** qPCR analysis was performed on the dissected ovaries of 6 and 13±1 days old females by using primers targeting the specific *Wolbachia* strains characterizing the two *Ae. albopictus* populations. Data were normalized by using DNA (2 ul of purified DNA per reaction). **B**: qPCR analysis was performed on bodies lacking ovaries of 6 and 13±1 days old females by using primers targeting the specific *Wolbachia* strains characterizing the two *Ae. albopictus* populations. Data were normalized by using the actin gene as host reference. Within each infection type and female age, different letters indicate statistically significant differences between treatments (One-way ANOVA followed by Tukey’s test: P<0.05).

When testing females without ovaries, the qPCR amplification results of *Wolbachia* (somatic fraction) did not reveal differences between treatments related to the titers of *w*AlbA, *w*AlbB, and *w*Pip *Wolbachia* when irradiated females had been tested at 6±1 days (Fig. 5B) (Kruskal Wallis test; *w*AlbA: χ^2^ = 1.85, df = 1, P = 0.17; *w*AlbB: χ^2^ = 0.32, df = 1; P = 0.57; *w*Pip: χ^2^ = 1.37, df = 1, P = 0.24). In contrast with the results obtained in the ovaries, the titer of *Wolbachia* in the extra-ovarian tissues exhibited an apparent age-related increase in irradiated females compared to the controls but differences between treatments were not significant (Kruskal Wallis test; *w*AlbA: χ^2^ = 2.17, df = 1, P = 0.14; *w*AlbB: χ^2^ = 2.52, df = 1, P = 0.11; *w*Pip: χ^2^ = 0.24, df = 1, P =0.62).

### FISH

The results of quantitative PCR were consistent with FISH images that revealed evident histological damage due to radiation and reduction in the mean size of the organs (Additional file 4: Figure S4), coupled with a markedly reduced *Wolbachia*-specific fluorescence intensity (Fig. 6; Additional files 5, 6, 7: Figures S5, S6, S7). Analysis of the *Wolbachia-*infected individuals revealed that fluorescent signals differed markedly between treatments (Fig. 6). In most of the ovaries dissected from S_ANG_ and AR*w*P females irradiated at 45 Gy, a weak fluorescent signal was observed (Fig. 6B, 6D; Additional files 6, 7: Figures S6, S7) compared to that in untreated controls (Fig. 6A, 6C; Additional files 8, 9: Figures S8, S9). In certain cases, the fluorescence was only localized in the ovarioles that possibly had not been degenerated by the irradiation (Fig. 6D, red circle). In contrast, the organs maintained their typical ovariole structure in untreated samples and the signal was abundant, specific, and confined within each oocyte (Fig. 6; Additional files 8, 9: Figures S8, S9).

**Figure 6.**
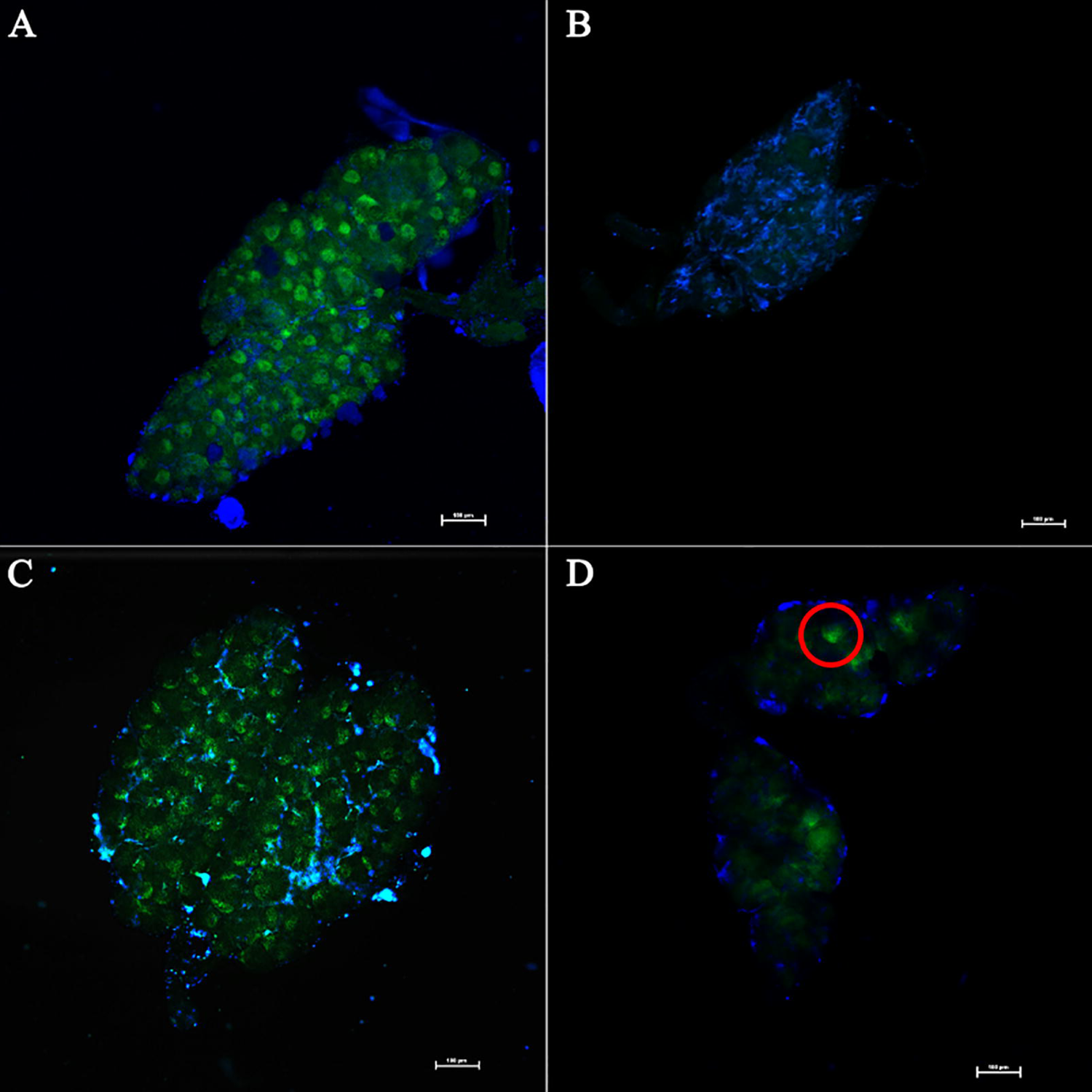
Fluorescence In Situ Hybridization of *Ae. albopictus* ovaries in irradiated females. A 45-Gy dose was applied to 36±4 hours old pupae of S_ANG_ and AR*w*P *Ae. albopictus*. Ovaries from sample individuals aged 13±1 days were then subjected to FISH analysis and compared with untreated counterparts. The distribution of *Wolbachia* is evidenced in green while blue stain is DAPI. **A**: Ovaries of untreated S_ANG_ females; **B**: Ovaries of irradiated S_ANG_ females **C**: Ovaries of untreated AR*w*P females; **D**: Ovaries of irradiated AR*w*P females.

## Discussion

Multiple blood-feeding is known to naturally occur in *Ae. albopictus* as not all of the ovarian follicles commence the gonotrophic cycle with a single blood meal [49]. Studies conducted on engorged *Ae. aegypti* females demonstrated that they were generally inhibited from host-seeking by the occurrence of distension-induced and oocyte-induced mechanisms of regulation [24, 50]; however, this inhibition does not apply to all individuals [49]. Results presented here demonstrate that the radiation doses generally used in the framework of SIT programs significantly attenuate this inhibition in both *Ae. albopictus* and *Ae. aegypti*. In fact, most females irradiated at 35 and 45 Gy fed several times a week, even if the highest number of bites was generally observed on the first day of the blood meal. Additionally, the dose of 28 Gy was found to be sufficient to induce a doubled biting rate (compared to the untreated controls). Tissue alterations in ovaries in response to irradiation might explain this phenomenon as the host-seeking behavior is interrupted through the release of hormones produced by the ovaries in the hemocoel during oogenesis [51, 52]. In virus-endemic areas, multiple blood feeding increases the likelihood of a female mosquito being infected by a suitable arbovirus and this possibility may significantly increase the chances of virus transmission to other hosts. Furthermore, the comparison of the survival curves of irradiated and untreated females revealed that the tested irradiation doses were not adequate to induce a marked decrease in life expectancy within the first 20 days in *Ae. albopictus*, which is an age that only a minority of adults are expected to reach in nature [53]. This suggests that irradiated *Ae. albopictus* females escaping the sexing procedure may have a stronger vectorial capacity than wild-type females.

The vectorial capacity of a species describes its potential to transmit a pathogen and is dependent on the ratio of mosquitoes to humans, the extrinsic incubation period of the parasite, the mosquitos’ rate of biting humans, and the survival of mosquito females [54–56]. SIT and SIT/IIT programs have the specific goal of reducing the vectorial capacity (together with the mosquito nuisance) via the suppression of a target population. In this context, the co-release of relatively small percentages of irradiated females with enhanced biting activity could be viewed as a negligible and transient side effect considering that the benefits are supposed to outweigh the potential risks. However, large scale releases of sterile males do not always adapt to the ever-decreasing wild population [19, 35] and if releases rely on high overflooding release ratios and a not perfect sexing, the increasing frequency of co-released irradiated females might represent an issue to be carefully evaluated in areas endemic for arboviral diseases.

Host-seeking is another behavioral trait that determines the biting rate and that could be affected by the radiation treatment. Previously, flight ability in radiation-sterilized males was not reported to be affected when compared to that in untreated males [9]. Our experiments testing the host-seeking ability in large enclosures confirm that flying ability is not affected by radiation doses that are capable of fully suppressing fecundity in females. Indeed, starved females irradiated at 45 Gy did not exhibit a significantly different ability to reach and bite the host compared to the controls, but when engorged, their behavior was anomalous and they continued to seek hosts, unlike the untreated females, leading to a marked increase in the mean number of bites per individual.

Overall, dose response curves related to fecundity and fertility were in agreement with the results of other studies [9] and verified the success of the radiation treatment. However, differently from previous reports [21], a dose of 28 Gy was inadequate to induce full sterilization in eggs. This result may indicate that this dose approaches the minimum threshold necessary to achieve full sterilization of *Ae. albopictus* females and that slight modifications of the experimental set up are sufficient to allow the viability of a small percentage of fertile eggs [19]. This issue also becomes evident when moving from laboratory conditions to large-scale operational conditions, and highlights the need for studies specific to the latter prior to conducting open field trials [22]. This is particularly necessary in relation to releases involving mosquito strains with altered *Wolbachia* infection types that have the ability to spread in the wild population through a Uni-CI pattern [15]. In these cases, a few partially fertile females could initiate a local population replacement with unpredictable consequences [20]. In this context, for safety purposes, adding a perfect sexing protocol to Uni-CI based IIT programs [17, 18, 35] could be preferable to combining IIT with SIT [19, 21, 22]. In fact, in order to avoid the release of fertile and potentially invasive females in a combined IIT/SIT approach, the required radiation doses may be so high as to seriously compromise male mating competitiveness [19] and may induce an enhanced biting behavior in the co-released females as evidenced in this study.

The evaluation of the vectorial capacity of a mosquito population also builds on the measurement of the vector competence which is mainly determined by genetic factors [57]. Due to the ability exhibited by *Wolbachia* of modulating these factors in infected mosquitoes, the decrease in density of these bacteria caused by the radiation treatment may introduce further concerns regarding the safe application of SIT in insect species infected by this bacterium when perfect sexing methods are not employed.

*Wolbachia*-mediated reduction of vector competence in mosquitoes has been suggested to depend on the bacterial titer and the specific strain of the bacterium [30, 58]. The density of *Wolbachia* in naturally infected *Ae. albopictus* is an individual feature that may vary substantially within a population [44] and that may be affected by the environmental conditions (temperature and food availability) under which the larvae develop [59–61]. Although distribution in somatic tissues has been observed in various cases, *Wolbachia* is mainly present within the germ line [62], and previous studies have already shown that the ovarian tissues of irradiated females are severely affected by the treatment [9, 21]. Bacteria are generally less sensitive to radiation than eukaryotes [63]; nevertheless, the density of *Wolbachia* is markedly affected by radiation doses employed for other insect species to achieve sterilization [28]. The results presented in this study highlighted an overall decrease in the density of *Wolbachia* in irradiated females, and this decrease was greatest when the pupae were irradiated at younger ages, and was positively correlated with female aging. This downward trend was common in all the studied *Wolbachia* strains (compared to the untreated controls).

Early irradiation is generally associated with a stronger efficiency of the sterilization treatment [43]; however, negative effects on fitness have been found to increase in younger treated pupae. For this reason, SIT programs are generally targeted at an intermediate pupal age to achieve full sterilization and to sufficiently preserve the fitness of adult males [9]. The damage experienced by the ovaries is consistent with complete sterilization that occurs when sufficiently high doses are employed. Our results highlighted that the destruction of these tissues—which are relatively sensitive to radiation due to intense mitotic activity [64]—seems to also affect the survival and/or the reproduction of the hosted *Wolbachia.* The endosymbiont may be involved—together with the host cells—in the mechanisms of apoptosis usually characterizing the oocytes subjected to irradiation [65]. However, further studies should be conducted to investigate this phenomenon and to determine why the reduction in the density of the bacterium becomes evident by qPCR only over a week after the treatment. This occurrence may be partly explained by a gradual clearance of the bacterial DNA corresponding to dead individuals or present in extracellular form [66] in the damaged ovarian tissues of irradiated females. Selection of an effective method to normalize the qPCR data allowed us to study an otherwise difficult-to-study phenomenon, as irradiation has previously been demonstrated to compromise the suitability of various common housekeeping genes when tissues are damaged [67]. A comparison of the actin gene copy number in the ovaries of irradiated and control individuals confirms this (Additional file2: Figure S2), and suggests the need for specific studies to identify the best candidate housekeeping genes when performing similar experiments. Investigating the levels of expression of specific *Wolbachia* genes, or host genes under *Wolbachia* control, may provide useful information to measure the activity of the bacteria, and as a consequence, to estimate their response to the irradiation treatment [68].

FISH supported the qPCR data related to ovaries, as demonstrated by the markedly reduced *Wolbachia*-specific fluorescence intensity associated with these tissues in irradiated females aged 13±1 days. This evidence reinforces the idea that the population of this endosymbiotic bacterium is severely affected by a 45-Gray radiation treatment, at least in this organ, and that qPCR is only partially capable of highlighting this phenomenon.

Overall, our results are insufficient to determine whether radiation treatment can be associated with a loss of specific *Wolbachia*-controlled biological activities such as pathogen interference. *Wolbachia* endosymbionts mainly exert their limiting action on arbovirus dissemination in the bodies of *Aedes* species by acting in the midgut [69], and similar to ovarian cells, midgut cells are characterized by intense mitotic activity and their structure and physiology have been previously shown to be affected by radiation [64, 70]. Therefore, *Wolbachia* and the related host-symbiont mechanisms of regulation may be affected by radiation in these tissues as well. However, this study does not provide evidence of such phenomena. Furthermore, as reported above, the detection of *Wolbachia* DNA does not necessarily indicate that the bacterium is alive or capable of fully exerting its effects on host physiology, because nucleic-acid-based analytical methods provide only limited information regarding the activities and physiological state of microorganisms in samples. These aspects can be detected retrospectively, but only after sufficient time has elapsed for the degradation and removal of DNA associated with inactivated cells [66, 71].

qPCR analysis specifically targeting the *Wolbachia* density in the gut of irradiated females may provide useful information to investigate bacterial population dynamics in these tissues. However, specific vector-competence studies will be necessary to ascertain whether a sterilizing radiation treatment leads to increased risk of virus acquisition or transmission per single bite. Coupled with the enhanced biting activity shown in the present study, an increased vector competence would boost further the vectorial capacity of female mosquitos. For such a study to be viable, oral infection trials should be conducted on mass-reared mosquito females after applying a radiation treatment at the doses necessary for large-scale operational programs, and all *Wolbachia*-infected vector species should be included [22].

In the case of programs based on the combination of SIT and IIT, investigation of the effects of irradiation on the induced level of CI in treated males should also be performed [31], because based on the results of the ovaries, a decrease in the titer of *Wolbachia* in the testes following the irradiation is a reasonably possible scenario.

## Conclusions

The results presented in this work stress the need for more thorough scientific investigations on the radiation biology of female mosquitoes. Such studies should be particularly opportune prior to conducting area-wide SIT or SIT/IIT programs that rely on imperfect sexing in areas endemic for arboviral diseases. Indeed, even a low percentage of contaminating mosquito females may result in the release of thousands of boosted vectors under large scale settings. Even if this safety issue should be negligible in the context of a successful population suppression program, conservatively, the effect of increasing frequencies of irradiated females in the target area should be thoroughly evaluated with the support of opportune models analyzing the epidemiological risks [55, 72, 73].

Measuring the vector competence of irradiated females that are infected with *Wolbachia* should be also opportune because the irradiation-induced decrease in the density of this bacterium may be consistent with effects on biological phenomena such as pathogen interference. Applying advanced and more efficient systems of sex separation capable of preventing the escape of females during release of sterile males [17, 18, 35, 74] would be sufficient to mitigate risks. Additionally, release protocols could use constant monitoring of the wild-type population to limit the number of mosquitoes released to a minimum threshold to guarantee efficacy [75]. Certainly, in the case of release programs that involve *Wolbachia* infections with pathogen interference phenotypes, the irradiation of pupae at young ages should be avoided because this treatment would maximize the biting activity and the *Wolbachia* depletion in adult females.

The data presented here may furnish useful cues to enhance the safety level of SIT-based control programs against *Aedes* mosquitoes and encourage a careful comparison between the various genetic control methods in search of the most efficient, sustainable, and safe strategy for mosquito vector control.

## List of Abbreviations

SIT: Sterile Insect Technique

CI: Cytoplasmic Incompatibility

Uni-CI: Unidirectional Cytoplasmic Incompatibility

IIT: Incompatible Insect Technique

qPCR: quantitative polymerase chain reaction

FISH: Fluorescence in situ hybridization

AR*w*P: *Aedes albopictus* population obtained through the replacement of the native

*w*AlbA-*w*AlbB: *Wolbachia* infection with *w*Pip *Wolbachia*

S_ANG_: Wild-type *Ae. albopictus* from Anguillara Sabazia (Rome)

LEU: large cellular polycarbonate experimental units

ANOVA: Analysis of viariance

## Declarations

### Ethics approval and consent to participate

Blood meals were provided via anesthetized mice in agreement with the Bioethics Committee for Animal Experimentation in Biomedical Research and in accordance with procedures approved by the ENEA Bioethical Committee according to the EU directive 2010/63/EU. The mice belonged to a colony housed at CR ENEA Casaccia and maintained for experimentation based on the authorization N. 80/2017-PR released (on February 2, 2017) by the Italian Ministry of Health.

Feeding female mosquitoes on the blood of human hosts (i.e., the Authors RM, EL, GL, and MC) during the experiments was also approved by the ENEA Bioethical Committee.

## Consent for publication

Two of the Authors (EL and MC) are present in Figure S1 and consent the publication of the image.

## Availability of data and materials

All relevant data are within the paper and its Supporting Information files. The datasets used and/or analysed during the current study are available from the corresponding author on reasonable request

## Authors’ contributions

RM and MC conceptualized the research. The methodology was planned by RM, EL, MC, AD, CD, and CP. RM, EL, GL, CD, CP, AD, and MC conducted the investigation. RM, EL, GF and MC organized and managed the datasets. RM, GF and MC conducted the statistical analysis of the data. MC and AS took care of funding acquisition. MC and CD supervised the experimentation. RM wrote the original draft of the manuscript. RM, MC, CD, EL, AS, GF, and GL contributed to writing, reviewing, and editing the manuscript.

## Acknowledgments

We thank Marta Piscitelli (Division for Health Protection Technologies, ENEA-Casaccia Research Center, Rome, Italy) as responsible of the facility for the housing and care of the mice used for blood feeding. We also thank Alessia Cappelli (School of Biosciences and Medical Veterinary, University of Camerino, MC, Italy) for her assistance with the FISH analysis and Alessia Fiore (Biotechnology and Agroindustry Division, ENEA-Casaccia Research Center, Rome, Italy) for her contribution to the normalization of the qPCR data through the measurement of the DNA content in the extracts from the ovarian tissues.

## Supplementary Information

Additional file 1: Figure S1. Schematic of host seeking behavior trials conducted under large enclosures

Additional file 2: Figure S2. Actin gene copies in the ovaries of *Ae. albopictus* females irradiated at 45 Gy in comparison with untreated counterparts

Additional file 3: Figure S3. Host seeking and biting behavior of irradiated S_ANG_ and AR*w*P *Ae. albopictus* under large enclosures compared to untreated controls. Biting proportions and average times to landing were compared between treatments within a 15-minute interval. **A:** Comparison between untreated and irradiated starved females aged 13±1 days; **B:** Comparison between irradiated and untreated females engorged females 48 hours after the engorgement (i.e., 15±1 days old). Two-way ANOVA demonstrated that the difference between treatments was statistically significant in the case of the engorged females (P < 0.05).

Additional file 4: Figure S4. Structural damage induced by irradiation at 45 Gy in the ovaries of 13±1 days old *Ae. albopictus* females in bright field**. A**: Ovaries of untreated S_ANG_ females; **B**: Ovaries of irradiated S_ANG_ females **C**: Ovaries of untreated AR*w*P females; **D**: Ovaries of irradiated AR*w*P females.

Additional file 5: Figure S5. FISH analysis of the ovaries of 13±1 days old *Ae. albopictus* belonging to a line (AR) cured from *Wolbachia* infection. **A** and **D**: DAPI stained; **B** and **E**: FITC stained; **C** and **F**: bright field. No specific green-fluorescent signal was detected in the aposymbiotic line.

Additional file 6: Figure S6. Additional images related to the FISH analysis of the ovaries of 13±1 days old S_ANG_ *Ae. albopictus* irradiated at 45 Gy. The distribution of *Wolbachia* is evidenced in green while blue stain is DAPI. **A** and **D**: DAPI stained; **B** and **E**: FITC stained; **C** and **F**: bright field. The green-fluorescent signal related to *Wolbachia* is weak and not homogeneously distributed.

Additional file 7: Figure S7. Additional images related to the FISH analysis of the ovaries of 13±1 days old AR*w*P *Ae. albopictus* irradiated at 45 Gy. The distribution of *Wolbachia* is evidenced in green while blue stain is DAPI. **A** and **D**: DAPI stained; **B** and **E**: FITC stained; **C** and **F**: bright field. The green-fluorescent signal related to *Wolbachia* is weak and not homogeneously distributed.

Additional file 8: Figure S8. Additional images related to the FISH analysis of the ovaries of 13±1 days old untreated AR*w*P *Ae. albopictus.* Blue stain is DAPI. **A** and **D**: DAPI stained; **B** and **E**: FITC stained; **C** and **F**: bright field. The green-fluorescent signal related to *Wolbachia* is strong and regularly distributed.

Additional file 9: Figure S9. Additional images related to the FISH analysis of the ovaries of 13±1 days old untreated S_ANG_ *Ae. albopictus*. Blue stain is DAPI. **A** and **D**: DAPI stained; **B** and **E**: FITC stained; **C** and **F**: bright field. The green-fluorescent signal related to *Wolbachia* is strong and regularly distributed.

*Article within a journal (no page numbers)*

Rohrmann S, Overvad K, Bueno-de-Mesquita HB, Jakobsen MU, Egeberg R, Tjønneland A, et al. Meat consumption and mortality - results from the European Prospective Investigation into Cancer and Nutrition. BMC Medicine. 2013;11:63.

*Article within a journal by DOI*

Slifka MK, Whitton JL. Clinical implications of dysregulated cytokine production. Dig J Mol Med. 2000; doi:10.1007/s801090000086.

*Article within a journal supplement*

Frumin AM, Nussbaum J, Esposito M. Functional asplenia: demonstration of splenic activity by bone marrow scan. Blood 1979;59 Suppl 1:26-32.

*Book chapter, or an article within a book*

Wyllie AH, Kerr JFR, Currie AR. Cell death: the significance of apoptosis. In: Bourne GH, Danielli JF, Jeon KW, editors. International review of cytology. London: Academic; 1980. p. 251-306.

*OnlineFirst chapter in a series (without a volume designation but with a DOI)*

Saito Y, Hyuga H. Rate equation approaches to amplification of enantiomeric excess and chiral symmetry breaking. Top Curr Chem. 2007. doi:10.1007/128_2006_108.

*Complete book, authored*

Blenkinsopp A, Paxton P. Symptoms in the pharmacy: a guide to the management of common illness. 3rd ed. Oxford: Blackwell Science; 1998.

